# Injury induced renal fibrosis promotes cystogenesis and cyst growth in adult mice with autosomal dominant polycystic kidney disease

**DOI:** 10.1101/2023.10.08.561382

**Authors:** Ming Wu, Dongping Chen, Yanzhe Wang, Pinglan Lin, Yanfang Bai, Yufeng Xing, Di Huang, Chaoyang Ye

## Abstract

Autosomal dominant polycystic kidney disease (ADPKD) is a life-long disease and caused by mutations in *PKD1* or *PKD2* gene. Fibrosis is a hallmark of chronic kidney disease and is positively correlated with renal cyst growth, however the role of fibrosis in ADPKD is still controversial. In this study, we established renal fibrosis by toxic or surgical injuries in adult mice, and *Pkd* gene was inactivated at different time point before or after renal injury according to the pattern of fibrosis progression in different injury models. Here we showed that renal injury before or after *Pkd* gene inactivation can both induce renal cysts in adult *Pkd1* or *Pkd2* mice, and the extent of cystic burden was tightly correlated with the baseline levels of fibrosis when three hits (injury and gene inactivation) occurred. Inactivation of *Pkd1* gene at the recovery stage after surgery induced less renal cysts in adult *Pkd1* mice. Enhanced renal fibrosis by repeated toxic injuries before gene inactivation accelerated renal cyst growth in *Pkd1* mice. We further showed that the rate of cyst formation at the early stage in adult *Pkd1* mice was positively correlated with the baseline levels of renal fibrosis. Finally, we showed that conditional knockout of *Ezh2* gene attenuated renal fibrosis and cyst growth in adult *Pkd1* mice with pre-existing renal fibrosis. We conclude that the fibrotic response after renal injury is a driving force for renal cyst formation and growth in adult kidneys and inhibition of renal fibrosis through targeting EZH2 might be new therapeutic strategy for adult ADPKD. Importantly, our study suggests that there is a time window for intervention upon acute kidney injury in adult ADPKD patients.

**Translational Statement:** Autosomal dominant polycystic kidney disease (ADPKD) is a life-long disease and caused by mutations in *PKD1* or *PKD2* gene. Fibrosis is a hallmark of chronic kidney disease and is positively correlated with renal cyst growth, however the role of renal fibrosis in ADPKD is controversial. In this study, we found that renal cysts were formed in adult *Pkd1* or *Pkd2* mice with established renal fibrosis induced by toxic or ischemia reperfusion injuries. Cyst formation or growth in adult ADPKD mice was tightly correlated with baseline levels of renal fibrosis after third hits. Enhanced renal fibrosis before *Pkd1* gene deletion in adult mice accelerated cyst growth. Inhibition of renal fibrosis through targeting EZH2 delayed cyst growth in adult ADPKD mice. Thus, renal fibrosis is a trigger of cyst formation and growth in adult ADPKD mice, and therapeutically targeting EZH2 might be new strategy to treat adult patients with ADPKD. Our study suggests that there is a time window for intervention upon acute kidney injury in adult ADPKD patients.

## Introduction

Autosomal dominant polycystic kidney disease (ADPKD) is the most common genetic kidney disease and one of the leading causes of end stage renal disease (ESRD) ^1,2^. It is caused by mutations in *PKD1* or *PKD2* gene and characterized by progressively enlargement of both kidneys which are filled with numerous cysts ^1,2^. Although renal cysts are mostly prenatally formed, ADPKD is an adult-onset disease, which leads to ESRD around 60 years of age in 70-75% of affected patients ^3–5^.

It was thought that renal cysts were developed in an autosomal recessive manner at cellular level, which means that germline (first hit) and somatic (second hit) mutations in both allele of *PKD* genes are required for cystogenesis ^6^. In line with this, only homozygous knockout of *Pkd1* gene induces renal cysts. Interestingly, immediate cyst formation is only induced in mice with *Pkd1* inactivation before postnatal day 13 ^7^. Adult inactivation of *Pkd1* gene results in renal cysts after 5 months ^7^. Ischemic or toxic renal injury accelerated cyst formation in adult *Pkd1* mice suggesting that a third hit (renal injury) is required for cystogenesis in matured kidneys ^8,9^. Injury induced kidney regeneration program constitutes activation of proliferating pathways, expression of acute injury kidney markers, induction of fibrotic responses, and activation of inflammatory responses, one of which could be the key mechanism triggering cyst formation in adult ADPKD kidneys ^6^.

Fibrosis is the hallmark of acute kidney injury (AKI) to chronic kidney disease (CKD) transition and is final common pathway of all kinds of CKD progressing to ESRD ^10^. Maladaptive repair of proximal tubules following recovery from severe and repeated episodes of AKI led to the development of renal interstitial fibrosis ^11^. Renal interstitial fibrosis is characterized by accumulation of extracellular matrix (ECM) proteins, over-expression of epithelial–mesenchymal transition (EMT) markers and activation of pro-fibrotic signaling pathways such as the TGFβ-Smad3 and Snail signaling pathway. Nearly every renal cyst has thickened basement membrane and ECM changes occur early in cyst development in human ADPKD samples ^2^. Extensive renal fibrosis is observed at the late stage of human ADPKD ^12^. In animals, knockout ECM component or inhibition of fibrosis related molecular pathways blocks renal cyst growth ^1,2^. All these data suggest that renal fibrosis is an essential component of cystogenesis and cyst development. However, in an orthologous *Pkd1* mouse model, collapsed cysts and reduced kidney weight were observed, which was accompanied by focal formation of fibrotic areas ^13^. Thus, renal fibrosis was thought to be simply a consequence of the pathology and may not be a driver of cyst development by some researchers in the field ^12^.

Animal models were established to study renal fibrosis according to the etiology of various CKD ^14^. Unilateral ischemia-reperfusion-induced injury (UIRI) and aristolochic acid I (AAI) induced renal fibrosis models are classic mouse models by mimicking the surgery or renal toxin induced nephropathy in humans ^14^.

In the current study, we established UIRI and AAN mouse models in adult ADPKD mice and inactivated *Pkd* genes at different time points according to the pattern of fibrosis progression in these two fibrosis models to study the role of renal fibrosis in adult ADPKD.

## Results

### Toxic injury induced fibrosis promoted the development of renal cysts in adult mice with *Pkd1* gene deletion

Mouse aristolochic acid nephropathy (AAN) model was established by intraperitoneally injection of aristolochic acid I (AAI, 5 mg/kg body weight). The time course expression of fibrotic markers (collagen-I, fibronectin, a-SMA, pSmad3) were assessed. Fibrotic markers in AAN kidneys were steadily increased from day 1 to day 21 (supl. Fig1). These data indicated that renal fibrosis was established at three weeks after AAI injection.

We next inactivated *Pkd1* gene at one week before AAI injection (early inactivation AAI-Pkd1 model) or at three weeks after AAI injection (late inactivation AAI-Pkd1 model) in 8 weeks adult *Pkd1^fl/fl^;Cre/Esr1^+^* mice (Figure 1A). Mice were sacrificed at 10 weeks after AAI injection (three hits occurred) in the early inactivation model or 10 weeks after *Pkd1* gene inactivation (three hits occurred) in the late inactivation model (Figure 1A). No renal cyst was observed in normal saline treated control *Pkd1^fl/fl^;Cre/Esr1^+^* mice and numerous small renal cysts were observed in early inactivated *Pkd1* mice as shown by HE staining (Figure 1B). Surprisingly, numerous big renal cysts were observed in late inactivated *Pkd1* mice (Figure 1B). Kidney weight was increased in early inactivated *Pkd1* mice as compared with normal saline treated control *Pkd1^fl/fl^;Cre/Esr1^+^* mice, which was further increased in the late inactivation model (Figure 1B). blood urine nitrogen (BUN) and serum creatinine (Scr) levels were only significantly increased in the late inactivation model as compared with normal saline treated control *Pkd1^fl/fl^;Cre/Esr1^+^* mice (Figure 1B).

**Fig. 1.**
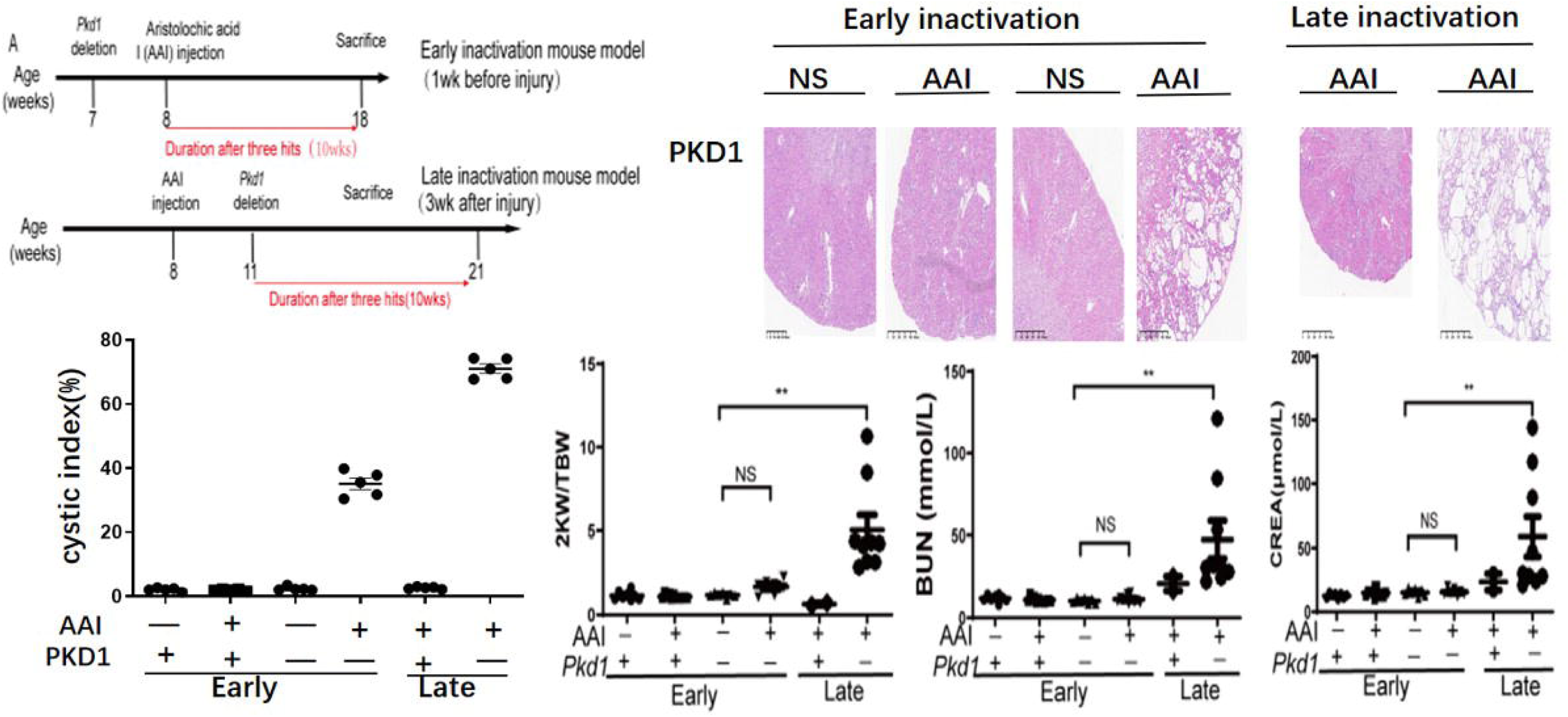
Toxic injury induced fibrosis promoted the development of renal cysts in adult mice with *Pkd1* gene deletion. (A) Schematic diagram of the experimental design *in vivo*. Renal injury was established by injection of aristolochic acid I (AAI) in male *Pkd1* mice at 8 weeks of age, and *Pkd1* gene was inactivated at one week before or three weeks after the kidney injury. Mice were sacrificed at 10 weeks after the three hits (kidney injury together with *Pkd1* gene deletion). (B-C) Renal cyst was assessed by hematoxylin-eosin (HE) staining and then quantified. Scale bar represents 100 μm. (D-F) Two kidney weight/total body weight (2KW/TBW) ratio, blood urine nitrogen (BUN), and serum creatinine (Scr) levels were assessed. N =6-8 in each experimental group and one representative of at least three independent experiments is shown. NS represents not significant. *p < 0.05. **p < 0.01. ***p < 0.001.

### Toxic injury induced fibrosis promoted the development of renal cysts in adult mice with *Pkd2* gene deletion

We further determined the role of fibrosis in *Pkd2* knockout (*Pkd2^fl/fl^;Cre/Esr1^+^*) mice with AAI induced fibrosis. Since cyst growth is quite aggressive in our *Pkd2* mouse model, we collected kidney samples at 7 weeks after three hits (*Pkd2* inactivation and toxic injury) (Figure 2A). Numerous small renal cysts were formed in early inactivated AAI-Pkd2 model as shown by HE staining (Figure 2B). Similar to *Pkd1* mouse model, numerous big renal cysts were observed in late inactivated AAI-Pkd2 model as compared to early inactivated AAI-Pkd2 model (Figure 2B). Kidney weight was increased in early inactivated *Pkd2* mice as compared with normal saline treated control *Pkd2^fl/fl^;Cre/Esr1^-^* mice, which was further increased in the late inactivation model (Figure 2B). BUN and Scr levels were only significantly increased in the late inactivation model as compared with normal saline treated control *Pkd2^fl/fl^;Cre/Esr1^-^* mice (Figure 2B).

**Fig. 2.**
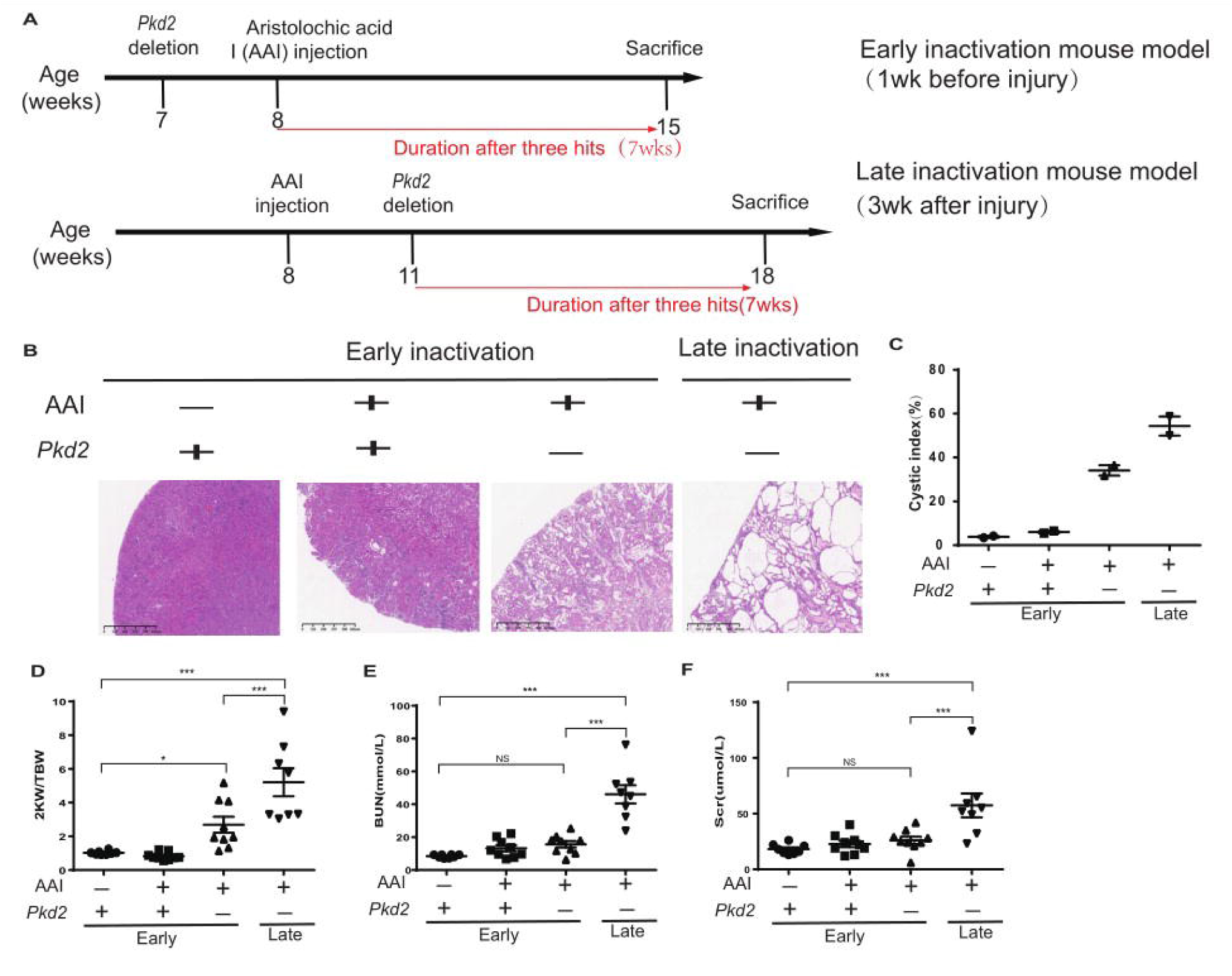
Toxic injury induced fibrosis promoted the development of renal cysts in adult mice with *Pkd2* gene deletion. (A) Schematic diagram of the experimental design *in vivo*. Renal injury was established by injection of aristolochic acid I (AAI) in male *Pkd2* mice at 8 weeks of age, and *Pkd2* gene was inactivated at one week before or three weeks after the kidney injury. Mice were sacrificed at 7 weeks after the three hits (kidney injury together with *Pkd2* gene deletion). (B-C) Renal cyst was assessed by hematoxylin-eosin (HE) staining and then quantified. Scale bar represents 100 μm. (D-F) Two kidney weight/total body weight (2KW/TBW) ratio, blood urine nitrogen (BUN), and serum creatinine (Scr) levels were assessed. N =6-8 in each experimental group and one representative of at least three independent experiments is shown. NS represents not significant. *p < 0.05. **p < 0.01. ***p < 0.001.

### Ischemic injury induced fibrosis promoted the development of renal cysts in adult mice with *Pkd1* gene deletion

Mouse model of renal unilateral ischemia-reperfusion-induced injury (UIRI) was established on the left kidney by 35 minutes of unilateral renal ischemia and followed by reperfusion for different time period. Fibrotic markers in UIRI kidneys were steadily increased from day 1 to day 7 and returned to the baseline after day 7 (supl. Fig2).

We inactivated *Pkd1* gene at one week before UIRI (early inactivation UIRI-Pkd1 model), at one week after UIRI (1wk-late inactivation UIRI-Pkd1 model) or at three week after UIRI (3wk-late inactivation UIRI-Pkd1 model) in 8 weeks adult *Pkd1^fl/fl^;Cre/Esr1^+^* mice (Figure 3A). Mice were sacrificed at 6 weeks after three hits (UIRI and *Pkd1* gene inactivation). No renal cysts was observed in normal saline treated control *Pkd1^fl/fl^;Cre/Esr1^+^*mice and a few focal renal cysts were observed in early inactivated UIRI-Pkd1 model (Figure 3B). Interestingly, numerous renal cysts were observed in the 1wk-late inactivation UIRI-Pkd1 model (Figure 3B). Surprisingly, very few renal cysts were observed in the 3wk-late inactivation UIRI-Pkd1 model (Figure 3B). Kidney weight was not increased in any group with *Pkd1* inactivation (Figure 3B).

**Fig. 3.**
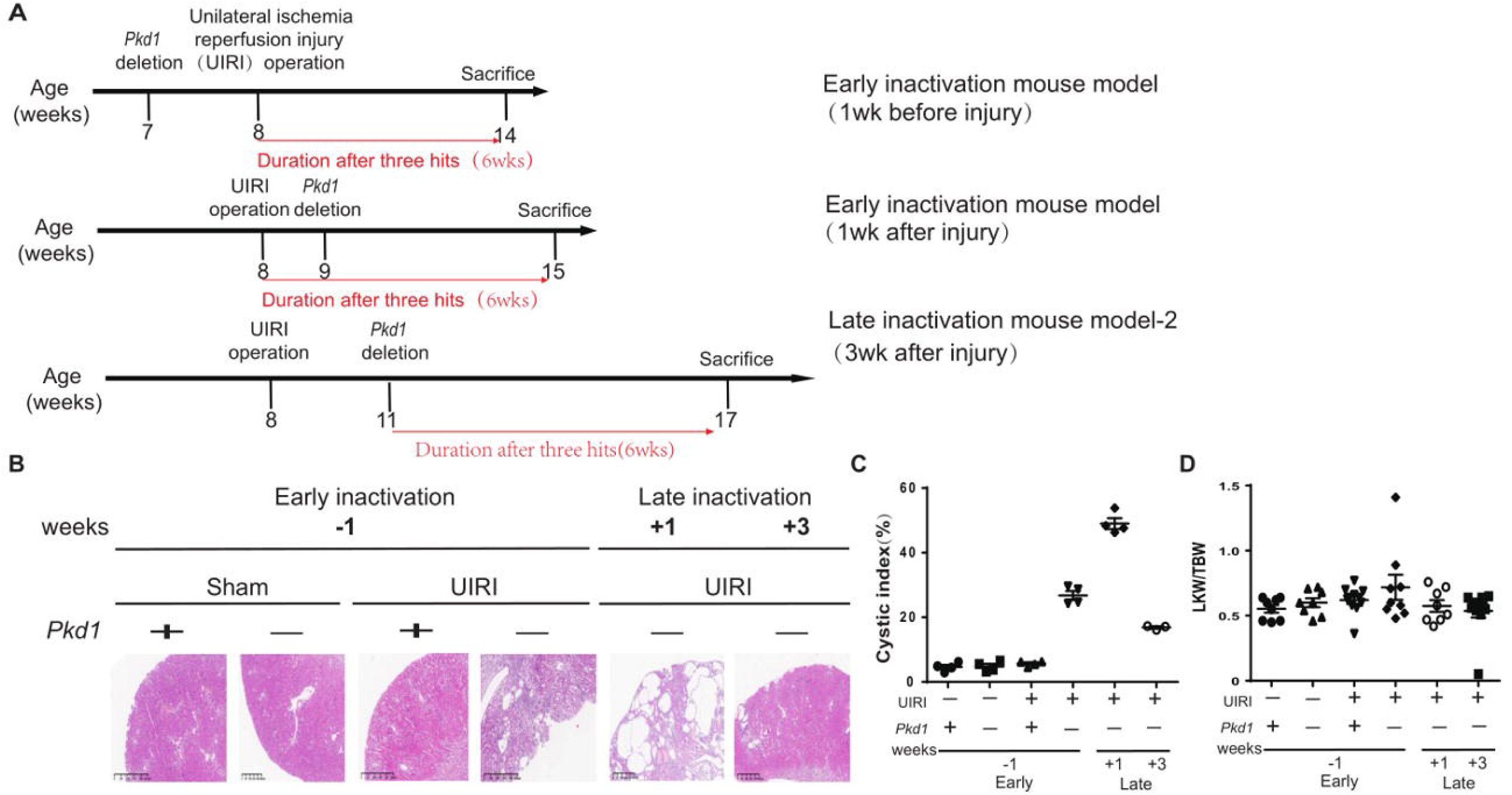
Ischemic injury induced fibrosis promoted the development of renal cysts in adult mice with *Pkd1* gene deletion. (A) Schematic diagram of the experimental design *in vivo*. Renal injury was established by unilateral ischemia-reperfusion injury (UIRI) on the left kidney in male *Pkd1* mice at 8 weeks of age, and *Pkd1* gene was inactivated at one week before or one week (1WK) or three weeks (3WK) after the kidney injury. Mice were sacrificed at 6 weeks after the three hits (kidney injury together with *Pkd1* gene deletion). (B-C) Renal cyst was assessed by hematoxylin-eosin (HE) staining and then quantified. Scale bar represents 100 μm. (D). Left kidney weight/total body weight (LKW/TBW) ratio was assessed. N =6-8 in each experimental group and one representative of at least three independent experiments is shown. NS represents not significant. *p < 0.05. **p < 0.01. ***p < 0.001.

### Enhanced fibrosis accelerates cyst growth in adult *Pkd1* mice

We enhanced the baseline levels of renal fibrosis by 3 times repeated injection of normal saline (NS) or low dose of AAI (2.5 mg/kg body weight) after initial one high dosage of AAI (5 mg/kg body weight) (supl. Fig3). As compared to repeated NS injected group, repeated injection of low dose AAI enhanced the expression of fibrotic markers in AAN kidneys (supl. Fig3).

After the initial high dose AAI injection in adult *Pkd1^fl/fl^;Cre/Esr1^+^* mice, normal saline (H+N) or low dose AAI (H+L) was repeatedly injected, which was followed by inactivation of *Pkd1* gene at three weeks after the initial high dose of AAI injection (Figure 4A). Kidneys were collected at seven weeks after *Pkd1* gene inactivation (Figure 4A). Cystic burden, kidney weight, BUN and Scr levels were significantly higher in AAI (H+L) model than that in AAI (H+N) model (Figure 4B).

**Fig. 4.**
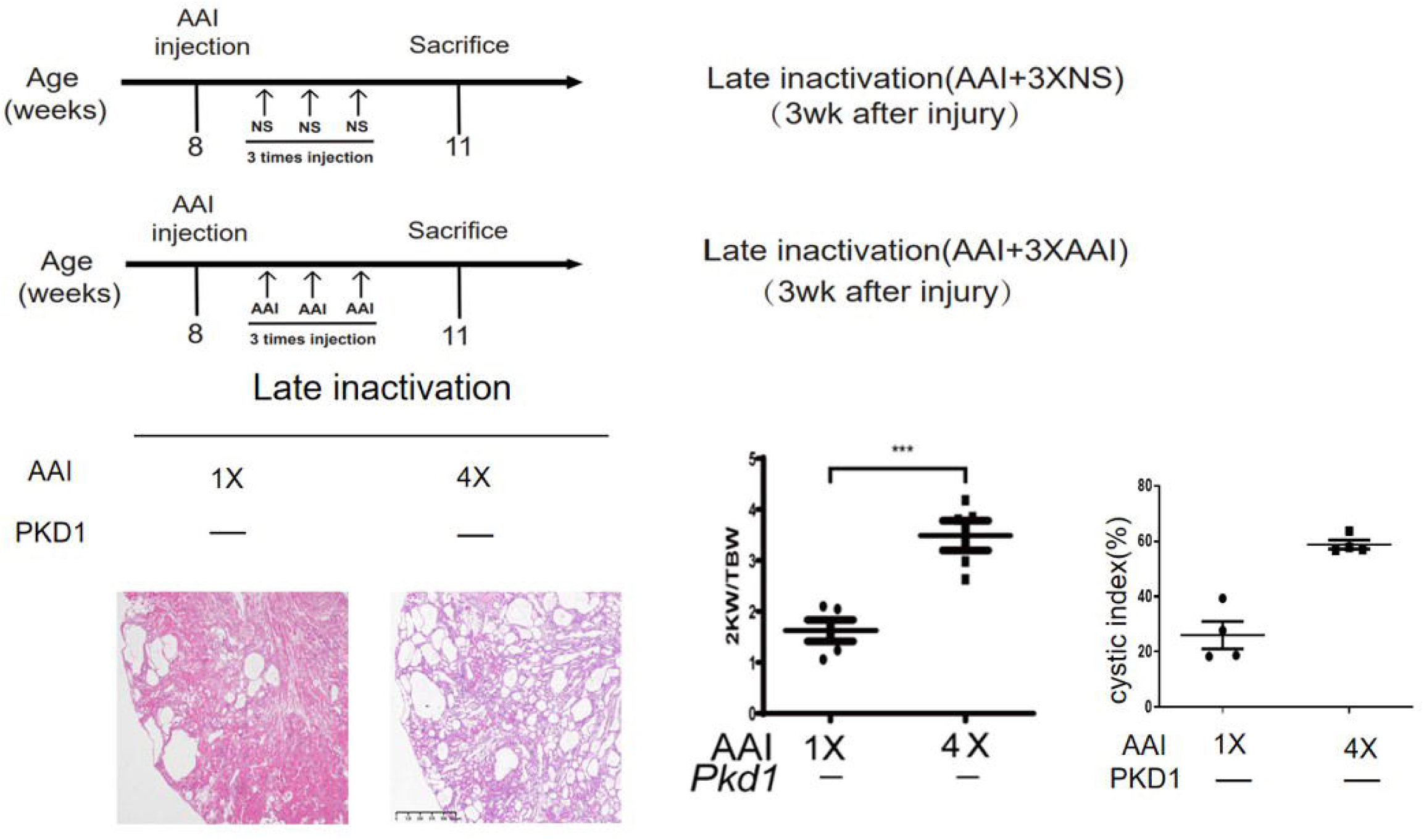
Enhanced fibrosis accelerates cyst growth in adult *Pkd1* mice. (A) Schematic diagram of the experimental design *in vivo*. Renal injury was established by injection of aristolochic acid I (AAI) in male *Pkd1* mice at 8 weeks of age, which was followed by three times injection of normal saline (NS) or AAI from one week after. *Pkd1* gene was inactivated at 11 weeks of age. Mice were sacrificed at 7 weeks after the three hits (kidney injury together with *Pkd1* gene deletion). (B-C) Renal cyst was assessed by hematoxylin-eosin (HE) staining and then quantified. Scale bar represents 100 μm. (D) Two kidney weight/total body weight (2KW/TBW) ratio, blood urine nitrogen (BUN), and serum creatinine (Scr) levels were assessed. N =6-8 in each experimental group and one representative of at least three independent experiments is shown. NS represents not significant. *p < 0.05. **p < 0.01. ***p < 0.001.

### Established renal fibrosis accelerates cystogenesis in adult *Pkd1* mice

Cystogenesis of *Pkd1* mice in the early stage in AAI early inactivation model and AAI late inactivation model was assessed (Figure 5A). In early inactivation AAI-PKD model, no cyst formation can be observed at day 21 after the third hit (*Pkd1* inactivation) (Figure 5B). In late inactivation AAI-PKD model, cyst formation can not be observed at day 10 after the third hit (AAI injection), however dilated tubules and few small cysts can be observed at day 21 (Figure 5B).

**Fig. 5.**
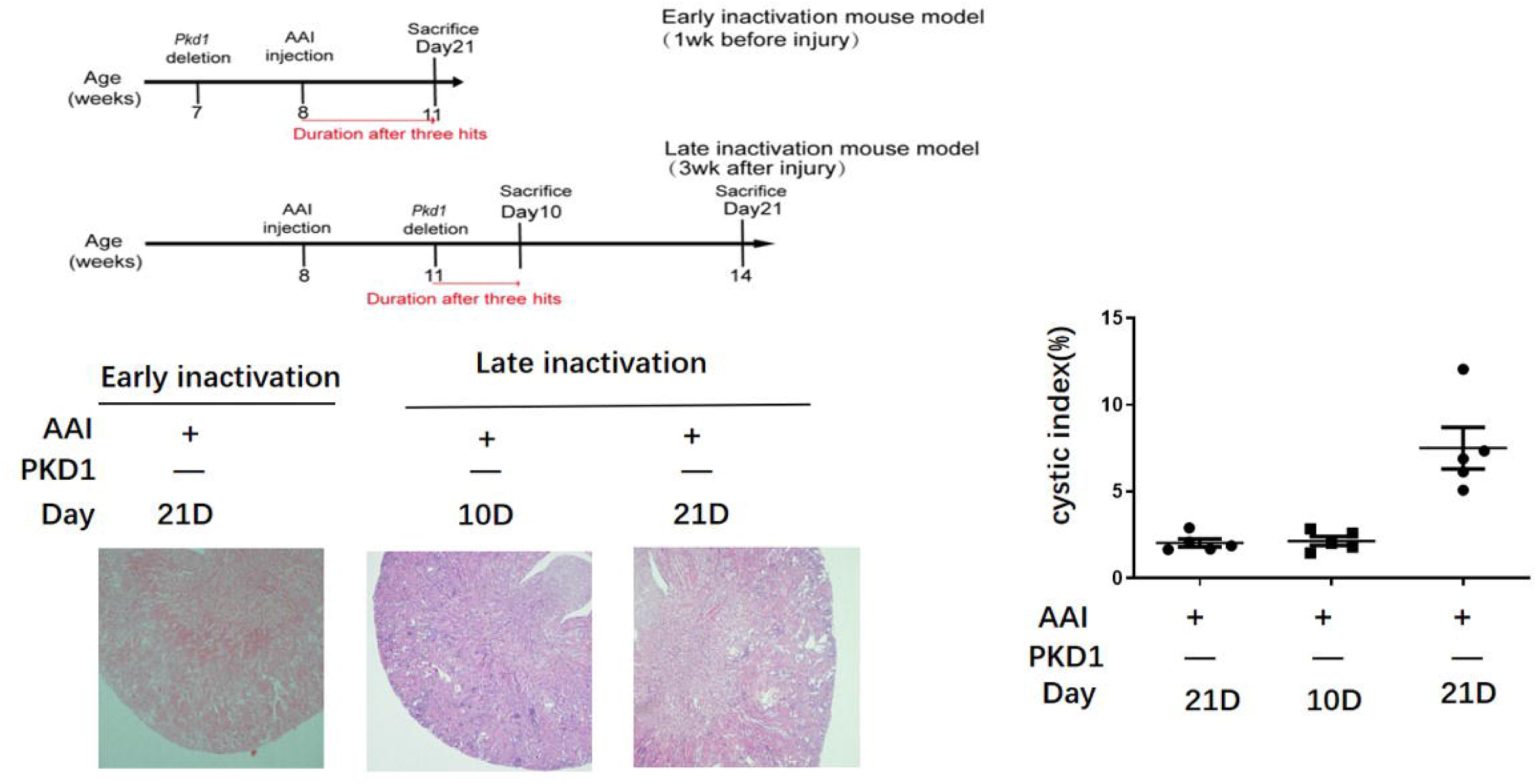
The rate of cyst formation at the early stage in adult Pkd1 mice with or without pre-established renal fibrosis. (A) Schematic diagram of the experimental design *in vivo*. Renal injury was established by injection of aristolochic acid I (AAI) in male *Pkd1* mice at 8 weeks of age, and *Pkd1* gene was inactivated at one week before or three weeks after the kidney injury. Mice were sacrificed at day 10 or day 21 after the three hits (kidney injury together with *Pkd1* gene deletion). (B-C) Renal cyst was assessed by hematoxylin-eosin (HE) staining and then quantified. Scale bar represents 100 μm. N =6-8 in each experimental group and one representative of at least three independent experiments is shown. NS represents not significant. *p < 0.05. **p < 0.01. ***p < 0.001.

### Conditional knockout of Ezh2 attenuated renal fibrosis and cyst growth in Pkd1 mice with established renal fibrosis

The epigenetic regulator EZH2 plays an important role in renal fibrosis ^15^. We hypothesized that inhibition of renal fibrosis through EZH2 retard disease progression in fibrosis triggered adult ADPKD. We first assessed the time course expression of EZH2 in the AAN mouse model. The expression of EZH2 is up-regulated in WT kidneys at 3 weeks after AAI injection (supl. Fig4).

Inactivation of *Ezh2* or *Pkd1* gene was performed in adult *Pkd1^fl/fl^;Cre/Esr1^+^* or *Ezh2 ^fl/fl^; Pkd1^fl/fl^;Cre/Esr1^+^* mice at three weeks after initial AAI injection (Figure 6A). Mice were sacrificed at 10 weeks after gene inactivation in the late inactivation model (Figure 6A). Cystic burden, kidney weight, BUN and Scr levels were significantly lower in *Ezh2 ^fl/fl^; Pkd1^fl/fl^;Cre/Esr1^+^* mice than that in *Pkd1^fl/fl^;Cre/Esr1^+^*mice (Figure 6B). Western blotting analysis showed that the expression of EZH2 and fibrotic markers were significantly lower in *Ezh2 ^fl/fl^; Pkd1^fl/fl^;Cre/Esr1^+^* mice than that in *Pkd1^fl/fl^;Cre/Esr1^+^* mice (Figure 6C).

**Fig. 6.**
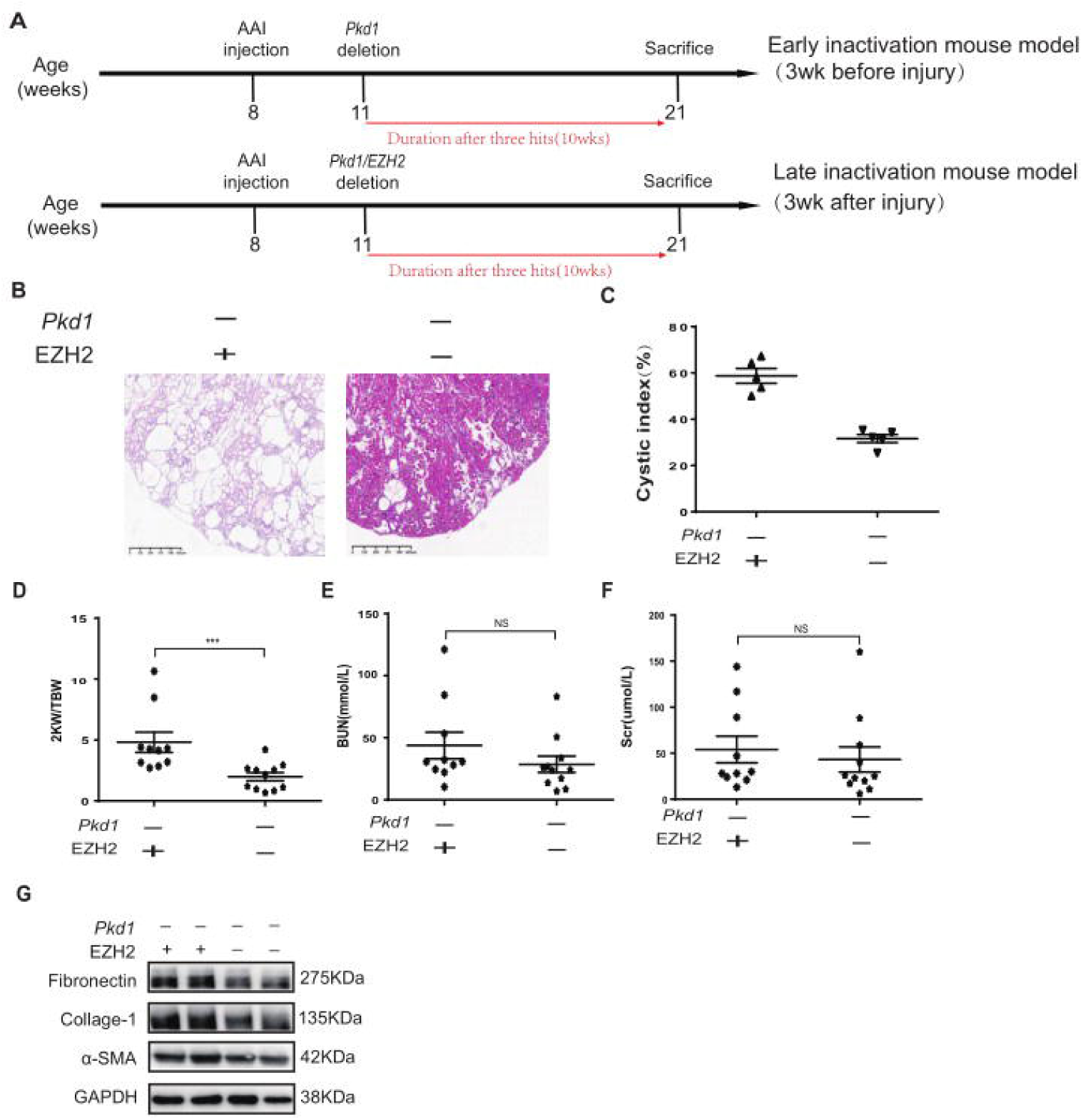
Conditional knockout of *Ezh2* gene attenuated renal fibrosis and cyst growth in adult *Pkd1* mice with established renal fibrosis. (A) Schematic diagram of the experimental design *in vivo*. Renal injury was established by injection of aristolochic acid I (AAI) in male mice at 8 weeks of age, and *Pkd1* and/or *Ezh2* gene was inactivated at three weeks after the kidney injury. Mice were sacrificed at 10 weeks after the three hits (kidney injury together with *Pkd1* gene deletion). (B-C) Renal cyst was assessed by hematoxylin-eosin (HE) staining and then quantified. Scale bar represents 100 μm. (D-F) Two kidney weight/total body weight (2KW/TBW) ratio, blood urine nitrogen (BUN), and serum creatinine (Scr) levels were assessed. (G). The expression of EZH2, collagen-I, and phosphorylated Smad3 (pSmad3) in mouse kidneys were analyzed by Western blotting and quantified. N =6-8 in each experimental group and one representative of at least three independent experiments is shown. NS represents not significant. *p < 0.05. **p < 0.01. ***p < 0.001.

## Discussions

In order to determine the role of fibrosis in ADPKD, we inactivated *Pkd1* or *Pkd2* gene according to the pattern of fibrosis progression in adult mice with toxic or surgery injuries. We found that renal cysts were developed in adult *Pkd1* or *Pkd2* mice with established renal fibrosis. Cyst formation or growth in adult ADPKD mice was tightly correlated with baseline levels of renal fibrosis after third hits (renal injury plus *Pkd* gene deletion). Enhanced renal fibrosis before *Pkd1* gene deletion in adult mice accelerated cyst growth. Inhibition of renal fibrosis through targeting EZH2 delayed cyst growth in adult ADPKD mice. Thus, renal fibrosis is a driver of cyst development in adult ADPKD mice.

Fibrosis is positively correlated with disease progression in human ADPKD ^12^. However, the role of fibrosis in ADPKD is controversial in animal studies ^1^. Reduced kidney weight and collapse of renal cysts were observed in an orthologous *Pkd1*^nl,nl^ mouse model after initial 4-week fast and massive cyst growth, and the areas of collapsed cysts were occupied with fibrotic tissues, suggesting that massive renal fibrosis does not support a further growth of enlarged cysts at the advanced stage of ADPKD when kidney size reaches a peak ^13^. However, a positive correlation of renal fibrosis with cyst growth was observed at the initial stage of cyst growth in *Pkd1*^nl,nl^ mice ^13^. Combined with our data, we suggest that renal fibrosis plays an important role at the early and middle stage of cyst development in adult ADPKD mice. ADPKD is a life-long disease, and the disease progression is accelerated around 50 years of age and progress to end-stage renal disease (ESRD) at a median age of 58 years ^5^. Thus, there is a large time window for therapeutic intervention in adult ADPKD patients to prevent or delay their progression into the advanced stage probably by managing the progression of renal fibrosis which could be induced by different etiologic causes in adult ADPKD. Importantly, our study suggests that there is also a short time window for intervention after each episode of acute kidney injury to prevent the growth of renal cysts in adult ADPKD by attenuating injury induced fibrotic responses.

The proximal tubule is the most vulnerable target after renal injury and hence is the focus of studies in the field of AKI and CKD ^1^^6,17^. Renal cysts are predominantly originated from collecting ducts in humans, and thus the proximal tubule is not the focus of ADPKD study so far.^18^ Interestingly, the injury of proximal tubule was observed in ADPKD kidneys, which is probably caused by the cyst expansion induce tubular obstruction ^19^. The AAN and UIRI mouse modes in our study are classic models to study renal fibrosis where the primary injury is taken place in the proximal tubule ^14,17^. Thus, our study suggests that a crosstalk may exist between injured proximal tubules and collecting ducts in ADPKD.

Partial EMT is a hallmark of renal fibrosis ^20^. Injured tubular cells undergo partial EMT, which will not migrate into interstitial areas but secret soluble factors to stimulate the growth and activation of interstitial fibroblasts ^20^. Our results indicate that there is a strong positive correlation of EMT and cyst growth, implying an EMT-mediated mechanism triggering cyst formation and growth.

Epigenetic regulation has been recently identified as an important mechanism involved in EMT and renal fibrosis ^21^. Previous studies showed that the epigenetic regulator EZH2 is up-regulated in fibrotic kidneys and pharmaceutical inhibition of EZH2 attenuates renal interstitial fibrosis in animals ^21^. The role of EZH2 in ADPKD is currently unknown. We hypothesized that inhibition of renal fibrosis by targeting EZH2 can retard ADPKD progression. Here, we showed that EZH2 expression is enhanced in AAI induced fibrotic kidneys. Deletion of EZH2 attenuated renal fibrosis and retarded disease progression in *Pkd1* mice.

In conclusion, fibrosis is a driving force of cystogenesis and cyst growth in adult mice with ADPKD. Therapeutic intervention of fibrosis is new strategy to delay disease progression in adult ADPKD.

## Materials and Methods

### Animals

Only male mice were used in animal studies. Wide type (WT) C57BL/6 mice (SPF grade, 20-25g) were purchased from Shanghai SLAC Laboratory Animal Co., Ltd. *Pkd1^fl/fl^* mice were obtained from Prof. Rudolf Wüthrich in University of Zurich, which were originally from the lab of Professor Gregory Germino in Johns Hopkins University School of Medicine ^22^. C57/BL6 *Pkd1^fl/fl^* mice were crossed with C57/BL6 tamoxifen-Cre (B6. Cg-Tg [Cre/Esr1]) 5Amc/J mice (stock 004682; Jackson Laboratories) as described previously ^23^. *Ezh2^fl/fl^* mice on a C57/B6 background were provided by Prof. Xi Wang from Capital Medical University and were further crossed with *Pkd1^f^ ^l/f^ ^l^;Cre/Esr1^+^* mice ^24^. Pkd2 conditional knockout mice with exon 3 deletion have cystic phenotype in the kidney ^25^. CRISPR/Cas9 technology was used to generate a mouse with loxP sites flanking exon 3 of the Pkd2 gene by Shanghai Model Organisms Center. *Pkd2^fl/fl^* were crossed with tamoxifen-Cre mice to generate *Pkd2^f^ ^l/f^ ^l^;Cre/Esr1^+^* mice. Animals were housed in the animal facility of Shanghai University of Traditional Chinese Medicine according to local regulations and guidelines. Animal experiments were approved by the ethic committee of Shanghai University of Traditional Chinese Medicine (PZSHUTCM18102604). Genotyping was performed for genetic animals and protocols will be provided upon request.

### Experimental Animal Protocol

To induce aristolochic acid nephropathy in mice, aristolochic acid I (AAI, A9451, Sigma; 5 mg/kg body weight) or normal saline (NS) was peritoneally injected in WT mice (25-30g). In another experiment, repeated low dose (2.5 mg/kg body weight) of AAI injection was followed by initial high dose of AAI injection. Mice were sacrificed at different time points (day 3 to 21) after initial high dose of AAI injection according to the experimental design, which is indicated in the result part. Kidney tissues were collected after sacrifice.

For the UIRI model, left renal pedicles were clamped for 35 minutes using microaneurysm clamps in male WT mice (20-25g). Mice were sacrificed at day 3, 7, 14 or 21 in a time-course experiment.

For conditional knockout mice, kidney injury (AAI injection or UIRI operation) was introduced at week 8 of age. Cre recombinase activity was induced by intraperitoneally injection of tamoxifen (50 mg kg-1 day-1, Apexbio, B5965) for three consecutive days at different time point according to the experimental design which is indicated in the result part.

### Cystic index analysis

Mouse kidneys were fixed and embedded in paraffin. Four-μm-thick sections of paraffin-embedded kidney tissue was sliced and subjected to hematoxylin and eosin (HE) staining. For cystic index analysis, 5 images were obtained for each section with the use of a microscope (Nikon 80i, Tokyo, Japan). Total area and cyst area measurement were evaluated using the ImageJ software. The cyst index was calculated as the ratio of cyst area to total area.

### Western Blotting Analysis

Cell or kidney protein was extracted by using lysis buffer bought from Beyotime Biotech (P0013, Nantong, China). The protein concentration was measured by using BCA Protein Assay Kit (P0012S, Beyotime Biotech, Nantong, China). Protein samples were dissolved in 5x SDS-PAGE loading buffer (P0015L, Beyotime Biotech, Nantong, China), which were further subjected to SDS-PAGE gel electrophoresis. After electrophoresis, proteins were electro-transferred to a polyvinylidene difluoride membrane (Merck Millipore, Darmstadt, Germany), which was incubated in the blocking buffer (5% non-fat milk, 20mM Tris-HCl, 150mMNaCl, PH=8.0, 0.01%Tween 20) for 1 hour at room temperature and was followed by incubation with anti-EZH2 (1:1000, 5246S, Cell Signaling Technology) antibody, anti-fibronectin (1:1000, ab23750, Abcam), anti-pSmad3 (1:1000, ET1609-41, HUABIO), anti-Collagen I (1:1000, ab260043, Abcam), anti-α-SMA (1:1000, ET1607-53, HUABIO), anti-N-cadherin (1:1000, sc-59887, Santa Cruz), anti-vimentin (1:1000, R1308-6, HUABIO), or anti-GAPDH (1:5000, 60004-1-lg, Proteintech) antibodies overnight at 4. Binding of the primary antibody was detected by an enhanced chemiluminescence method (SuperSignal™ West Femto, 34094, Thermo Fisher Scientific) using horseradish peroxidase-conjugated secondary antibodies (goat anti-rabbit IgG, 1:1000, A0208, Beyotime or goat anti-mouse IgG, 1:1000, A0216, Beyotime).

### Statistical Analysis

Statistical analyses were performed by one-way ANOVA followed by a Dunnett’s multiple comparisons test using GraphPad Prism version 5.0 (GraphPad, San Diego, CA). All data are expressed as means ± SD, and P < 0.05 was considered as statistically significant.

## Supplemental figure legends

Fig. S1. The time course expression of renal fibrosis in mouse kidneys with aristolochic acid induced nephropathy.

Mouse aristolochic acid nephropathy (AAN) model was established by intraperitoneally injection of aristolochic acid I (AAI, 5 mg/kg body weight). The expression of collagen-I, phosphorylated Smad3 (pSmad3) in normal saline (NS) or AAN kidneys were analyzed by Western blotting and quantified. N =6-8 in each experimental group and one representative of at least three independent experiments is shown. NS represents not significant. *p < 0.05. **p < 0.01. ***p < 0.001.

Fig. S2. The time course expression of renal fibrosis in mouse kidneys with unilateral ischemia-reperfusion injury.

Mouse model of renal unilateral ischemia-reperfusion-induced injury (UIRI) was established on the left kidney by 35 min of unilateral renal ischemia in WT mice and followed by reperfusion for different time. The expression of collagen-I, phosphorylated Smad3 (pSmad3) in sham or UIRI kidneys were analyzed by Western blotting and quantified. N =6-8 in each experimental group and one representative of at least three independent experiments is shown. NS represents not significant. *p < 0.05. **p < 0.01. ***p < 0.001.

Fig. S3. Repeated toxic injuries enhanced renal fibrosis.

(A) Schematic diagram of the experimental design *in vivo*. WT mice were repeated injected with NS or low dose of AAI (2.5 mg/kg body weight) at one week after initial one high dosage of AAI (5 mg/kg body weight). (B) The expression of collagen-I, phosphorylated Smad3 (pSmad3) in UIRI kidneys were analyzed by Western blotting and quantified. N =6-8 in each experimental group and one representative of at least three independent experiments is shown. NS represents not significant. *p < 0.05. **p < 0.01. ***p < 0.001.

Fig. S4. The time course expression of EZH2 in mouse kidneys with aristolochic acid induced nephropathy.

Mouse aristolochic acid nephropathy (AAN) model was established by intraperitoneally injection of aristolochic acid I (AAI, 5 mg/kg body weight). The expression of EZH2 in normal saline (NS) or AAN kidneys were analyzed by Western blotting and quantified. N =6-8 in each experimental group and one representative of at least three independent experiments is shown. NS represents not significant. *p < 0.05. **p < 0.01. ***p < 0.001.

## References

1. Fragiadaki, M., Macleod, F.M. & Ong, A.C.M. The Controversial Role of Fibrosis in Autosomal Dominant Polycystic Kidney Disease. Int J Mol Sci 21(2020).

2. Zhang, Y., Reif, G. & Wallace, D.P. Extracellular matrix, integrins, and focal adhesion signaling in polycystic kidney disease. Cell Signal 72, 109646 (2020).

3. Bergmann, C., et al. Polycystic kidney disease. Nat Rev Dis Primers 4, 50 (2018).

4. Reiterova, J. & Tesar, V. Autosomal Dominant Polycystic Kidney Disease: From Pathophysiology of Cystogenesis to Advances in the Treatment. Int J Mol Sci 23(2022).

5. Gansevoort, R.T., et al. Recommendations for the use of tolvaptan in autosomal dominant polycystic kidney disease: a position statement on behalf of the ERA-EDTA Working Groups on Inherited Kidney Disorders and European Renal Best Practice. Nephrol Dial Transplant 31, 337–348 (2016).

6. Weimbs, T. Third-hit signaling in renal cyst formation. J Am Soc Nephrol 22, 793–795 (2011).

7. Piontek, K., Menezes, L.F., Garcia-Gonzalez, M.A., Huso, D.L. & Germino, G.G. A critical developmental switch defines the kinetics of kidney cyst formation after loss of Pkd1. Nat Med 13, 1490–1495 (2007).

8. Takakura, A., et al. Renal injury is a third hit promoting rapid development of adult polycystic kidney disease. Hum Mol Genet 18, 2523–2531 (2009).

9. Happe, H., et al. Toxic tubular injury in kidneys from Pkd1-deletion mice accelerates cystogenesis accompanied by dysregulated planar cell polarity and canonical Wnt signaling pathways. Hum Mol Genet 18, 2532–2542 (2009).

10. Panizo, S., et al. Fibrosis in Chronic Kidney Disease: Pathogenesis and Consequences. Int J Mol Sci 22(2021).

11. Yu, S.M. & Bonventre, J.V. Acute kidney injury and maladaptive tubular repair leading to renal fibrosis. Curr Opin Nephrol Hypertens 29, 310–318 (2020).

12. Song, C.J., Zimmerman, K.A., Henke, S.J. & Yoder, B.K. Inflammation and Fibrosis in Polycystic Kidney Disease. Results Probl Cell Differ 60, 323–344 (2017).

13. Happe, H., et al. Cyst expansion and regression in a mouse model of polycystic kidney disease. Kidney Int 83, 1099–1108 (2013).

14. Fu, Y., et al. Rodent models of AKI-CKD transition. Am J Physiol Renal Physiol 315, F1098–F1106 (2018).

15. Zhou, X., et al. Enhancer of Zeste Homolog 2 Inhibition Attenuates Renal Fibrosis by Maintaining Smad7 and Phosphatase and Tensin Homolog Expression. J Am Soc Nephrol 27, 2092–2108 (2016).

16. Chevalier, R.L. The proximal tubule is the primary target of injury and progression of kidney disease: role of the glomerulotubular junction. Am J Physiol Renal Physiol 311, F145–161 (2016).

17. Gewin, L.S. Renal fibrosis: Primacy of the proximal tubule. Matrix Biol 68-69, 248–262 (2018).

18. Grantham, J.J. Rationale for early treatment of polycystic kidney disease. Pediatr Nephrol 30, 1053–1062 (2015).

19. Galarreta, C.I., et al. Tubular obstruction leads to progressive proximal tubular injury and atubular glomeruli in polycystic kidney disease. Am J Pathol 184, 1957–1966 (2014).

20. Sheng, L. & Zhuang, S. New Insights Into the Role and Mechanism of Partial Epithelial-Mesenchymal Transition in Kidney Fibrosis. Front Physiol 11, 569322 (2020).

21. Li, T., Yu, C. & Zhuang, S. Histone Methyltransferase EZH2: A Potential Therapeutic Target for Kidney Diseases. Front Physiol 12, 640700 (2021).

22. Su, Z., et al. Excessive activation of the alternative complement pathway in autosomal dominant polycystic kidney disease. J Intern Med 276, 470–485 (2014).

23. Sun, Y., et al. Activation of P-TEFb by cAMP-PKA signaling in autosomal dominant polycystic kidney disease. Sci Adv 5, eaaw3593 (2019).

24. Yin, J., et al. Ezh2 regulates differentiation and function of natural killer cells through histone methyltransferase activity. Proceedings of the National Academy of Sciences of the United States of America 112, 15988–15993 (2015).

25. Kim, I., et al. Conditional mutation of Pkd2 causes cystogenesis and upregulates beta-catenin. J Am Soc Nephrol 20, 2556–2569 (2009).

