## Supplementary figures and images for "Injury induced renal fibrosis promotes cystogenesis and cyst growth in adult mice with autosomal dominant polycystic kidney disease"

### Supplemental Figure-1

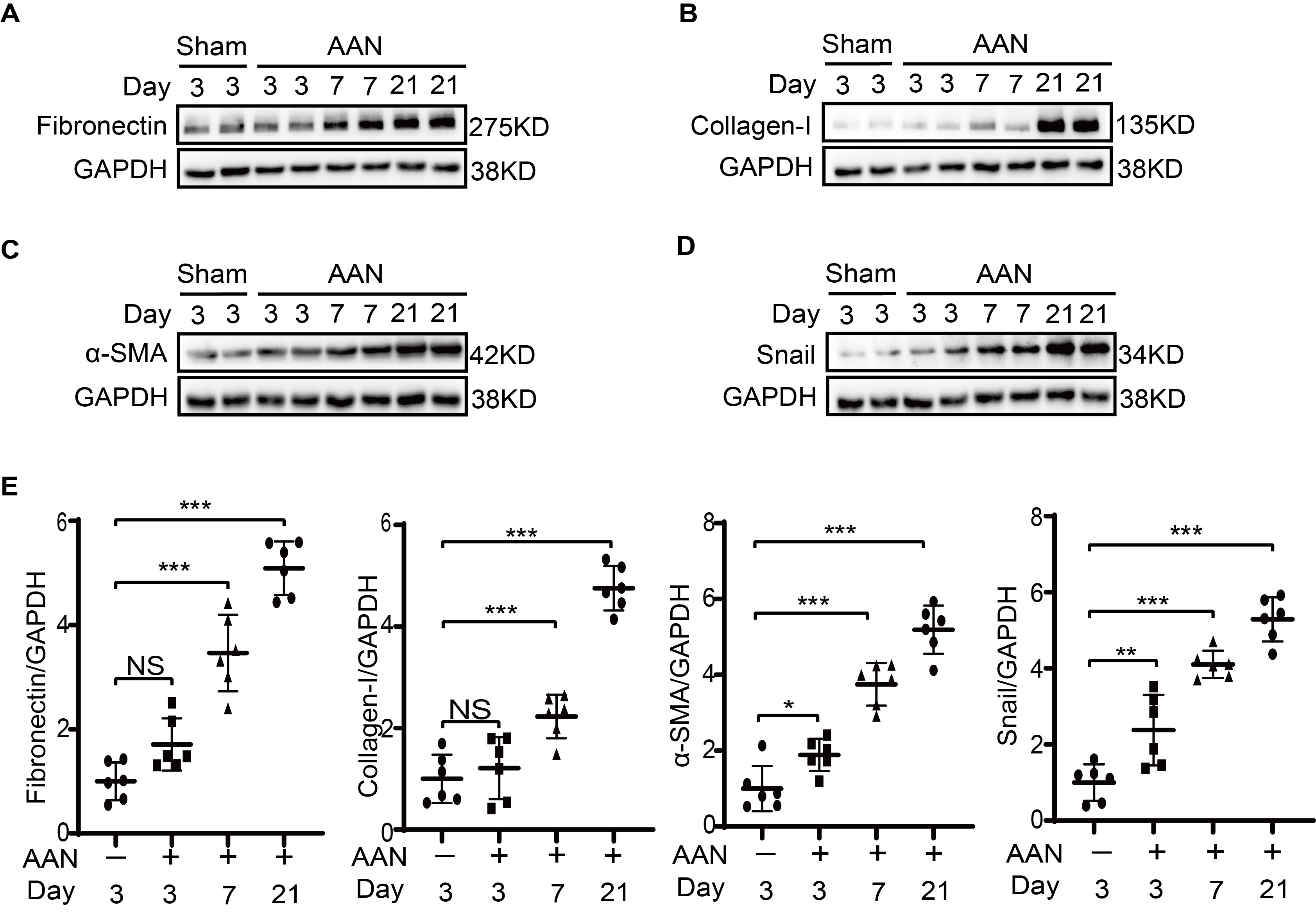

### Supplemental Figure-2

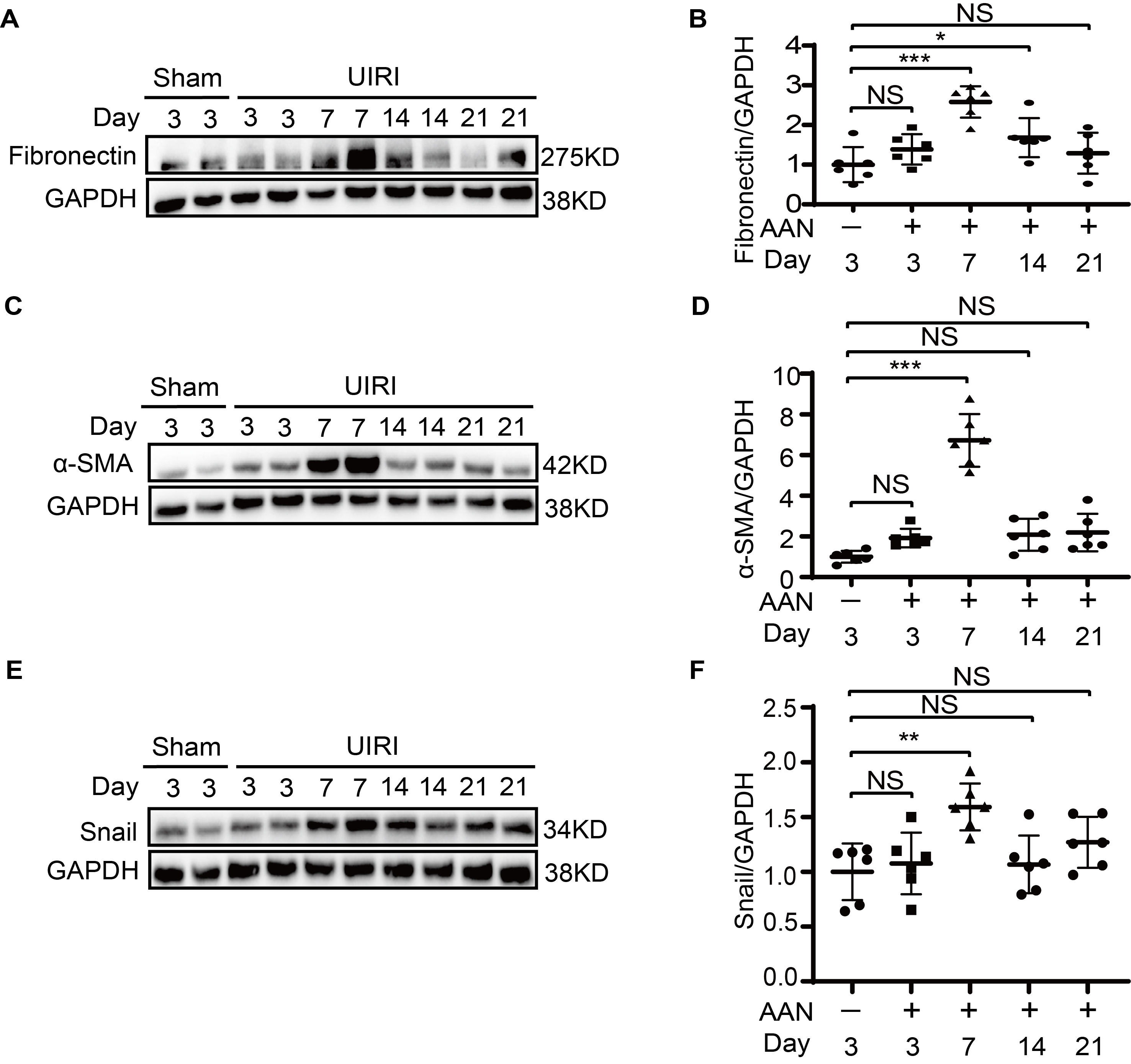

### Supplemental Figure-3

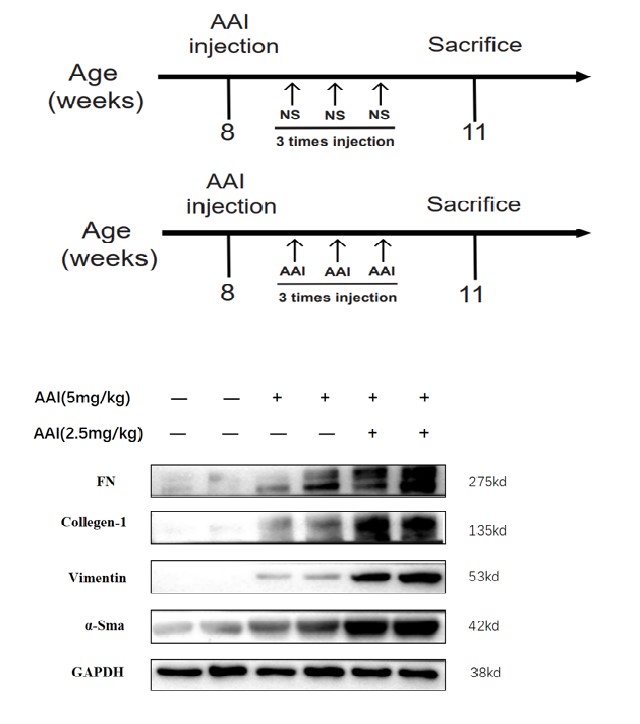

### Supplemental Figure-4

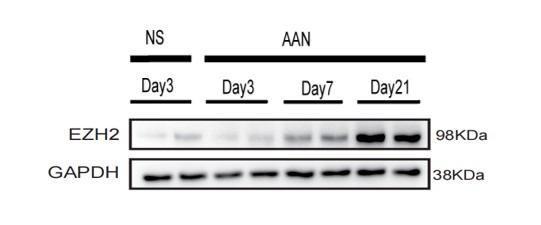
